# Variability in mosquito community composition associated with estuarine wetlands in Northern NSW, Australia

**DOI:** 10.1101/2021.12.13.472482

**Authors:** Cameron Ewart Webb, Jayne Hanford, Michael Bald, Scott Roberts

**Affiliations:** Marie Bashir Institute for Infectious Diseases and Biosecurity, The University of Sydney, NSW 2006, Australia; Medical Entomology, NSW Health Pathology, Westmead Hospital, Westmead, NSW 2145, Australia; School of Life and Environmental Sciences, The University of Sydney, Sydney, NSW 2000, Australia; Ardill Payne and Partners, Ballina NSW 2478, Australia

**Keywords:** *Aedes vigilax*, *Verrallina funerea*, *Culex sitiens*, mangroves, saltmarsh

## Abstract

The Northern Rivers region of NSW, Australia, is well documented as being impacted by nuisance-biting mosquitoes and mosquito-borne disease. Mosquitoes of greatest concern are those associated with estuarine and brackish water habitats associated with coastal wetlands and understanding the spatial variability in abundance and diversity will assist the assessment of risk and inform surveillance and control programs. Adult mosquito populations were sampled, using carbon dioxide baited traps, at four locations within the Richmond River estuary at Ballina, NSW, Australia, during January and February 2021. Concomitant sampling of habitats for immature mosquitoes was also undertaken. A total of 16,467 mosquitoes was collected at all sites across two sampling periods with the most abundant mosquitoes, *Verrallina funerea, Aedes vigilax*, and *Culex sitiens*, those typically associated with estuarine environments. *Culex annulirostris*, a mosquito associated with freshwater habitats, and *Aedes notoscriptus*, a mosquito associated with water-holding containers, were also commonly collected. The mosquito communities differed, in relative abundance and species richness, between the four locations. The result highlighted the need for authorities to understand the variability in productivity of potential mosquito habitats, beyond those determinants associated with vegetation communities alone, when assessing suitable locations of mosquito surveillance and integrated mosquito management.

## Introduction

Australia has a diverse, and often abundant, mosquito fauna (Webb et al. 2016). Mosquito-borne disease is a concern for health authorities where Ross River virus (RRV), and other arboviruses, pose a risk to human health (Claflin and Webb 2015). Approximately 5,000 cases of human disease resulting from RRV infection is reported annually across Australia (National Notifiable Diseases Surveillance Scheme 2021). Notwithstanding the public health concerns associated with mosquitoes, the impact of nuisance biting insects can also significantly impact the community, especially those living in close proximity to local wetland and bushland areas (Ratnayake et al. 2006). While there are ecological sustainable management programs in place in many regions (Russell and Kay 2008; Carlson et al. 2019), they assist with minimising nuisance-biting impacts but may not necessarily mitigate the public health risks associated with mosquitoes (Tomerini et al. 2011).

The Northern Rivers region of NSW experiences high levels of mosquito-borne disease activity on an annual basis, including occasional significant epidemics (Jansen et al 2020). For the past 20 years, among Northern NSW local health district residents, there has been an average of 162 confirmed and probable cases of disease caused by RRV reported to local authorities with a peak in activity during the 2019-2020 season of 420 cases (NSW Health Notifiable Conditions Information Management System (NCIMS), Communicable Diseases Branch and Centre for Epidemiology and Evidence, NSW Health 2021).While mosquitoes associated with a range of habitats within the region have the potential to impact the local community (Webb et al. 2016), the mosquitoes of greatest concern are those associated with the major estuaries where saltmarsh, mangrove, she-oak and coastal swamp forest habitats provide suitable conditions for mosquitoes such as *Aedes vigilax* Skuse (Diptera: Culicidae) and *Verrallina funerea* (Diptera: Culicidae). Mosquitoes found in freshwater wetlands (e.g. *Anopheles annulipes* (Diptera: Culicidae), *Aedes procax* (Diptera: Culicidae), *Coquillettidia linealis* (Diptera: Culicidae), *Culex annulirostris* (Diptera: Culicidae) and *Mansonia uniformis* (Diptera: Culicidae)), may also a pose a pest and public health threat.

One of the most active local authorities within the region undertaking management of mosquito impacts is Ballina Shire Council (BSC) who have participated in the NSW Health Arbovirus Surveillance and Mosquito Monitoring Program (NSW Health) for over 20 years (Doggett et al. 2009). While there are no local broadscale mosquito control programs within the local area, in response to the risk posed by mosquitoes, BSC have a number of initiatives that are designed to raise awareness of the risks associated with mosquitoes and promote safe and effective mosquito bite avoidance activity within the community and have also sought to guide planning decisions in the region where the pest and public health risks of mosquitoes is considered important (Dale et al. 2008; Dwyer et al. 2016; Carlson et al 2019; Ballina Shire Council 2021).

Critical to understanding the spatial and temporal variability in the pest and public health risks associated with mosquitoes is to determine their close associations with local habitats (Dale and Knight 2008). However, reliance on simplistic links between vegetation communities and the associated abundance and diversity of mosquito fauna may result in misleading indications of those risks (Claflin and Webb 2017). To better understand the spatial variability in abundance and diversity of mosquitoes associated with estuarine wetlands of the BSC region, adult and immature mosquito populations were sampled following environmental events, specifically rainfall and tidal inundation, known to drive activity of mosquitoes associated with these habitats.

## Methods

Mosquito populations were monitored during two periods; 13-15 January 2021 and 11-14 February 2021 at five locations in the local area (River Oaks Estate Conservation Area, Fishery Creek, Little Fishery Creek, North Creek, and Ballina Heights) (Fig.1). Four of these locations were situated within the tidally influenced North Creek estuary while the site a Ballina Heights was included as a reference location. The frequency of mosquito sampling was determined by prevailing environmental conditions and their potential impact on estuarine mosquito habitats (i.e. approximately two weeks following habitat inundation by tides and/or rainfall). Adult mosquitoes were collected using Encephalitis Virus Surveillance (EVS) traps (Rohe and Fall 1979). These traps use carbon dioxide to attract host-seeking female mosquitoes and are commonly used for monitoring mosquito populations of pest and public health concern in Australia (Webb et al 2016). Traps were set in the late afternoon and collected the following morning. Mosquito specimens were returned to the laboratory for identification according to the taxonomic keys of Russell (1993) and pictorial guides of Webb et al (2016) with the species richness and abundance of collections recorded according to sampling location and collection date.

**Figure 1.**
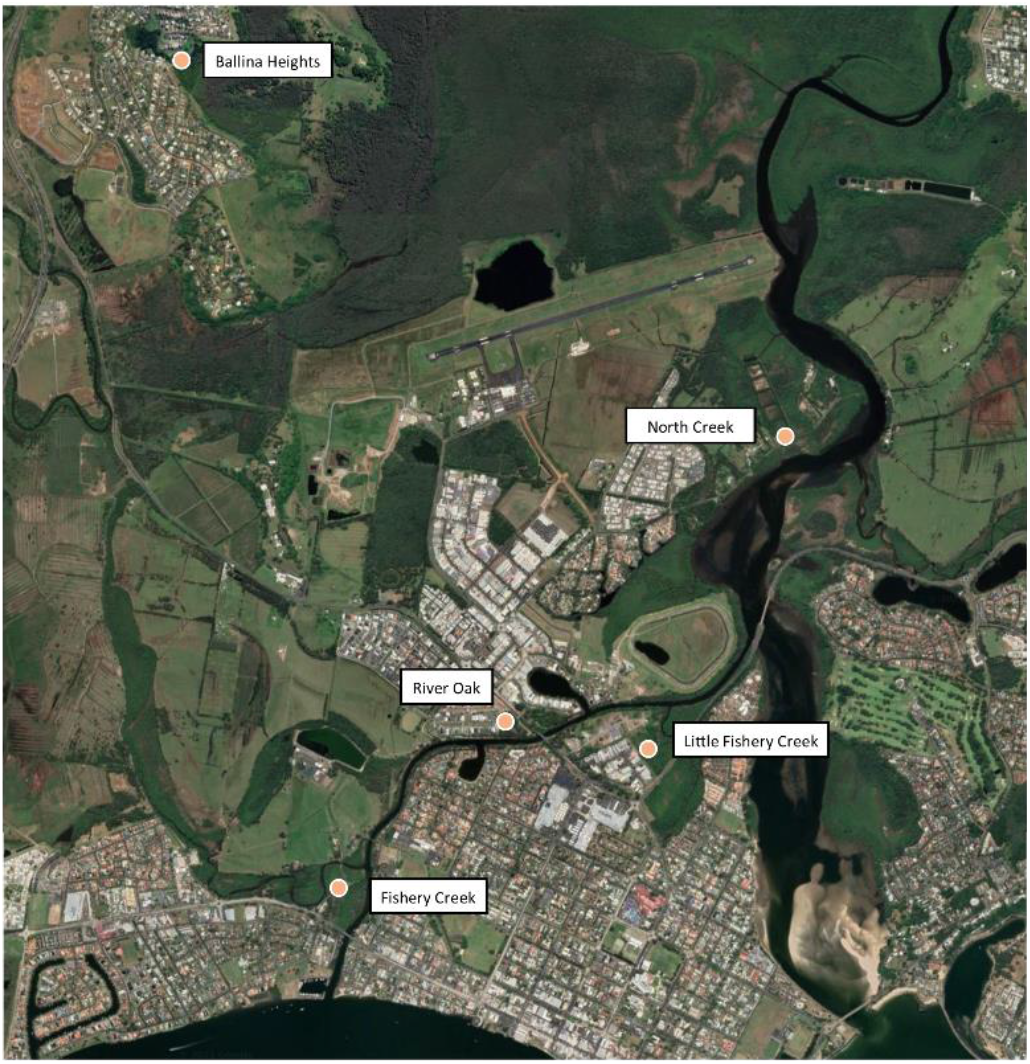
Overview of five study locations where mosquito sampling was undertaken to determine relative mosquito abundance associated with the River Oaks Estate Conservation Area, January-February 2021.

Mean daily maximum and minimum temperatures as well as monthly rainfall was provided by the Bureau of Meteorology (Ballina Airport AWS; Station ID 058198). Across the three month period, December 2020 through to February 2021, mean maximum (27.7°C) and mean minimum (20.1°C) daily temperatures varied marginally from long term average conditions of mean maximum (28.0°C) and minimum (19.3°C) daily temperatures. Over the same three month period, there was approximately 830mm of rainfall reported compared to the long term average of 516mm. Prior to the two survey periods, tides of over 1.8m were recorded in the Richmond River (Missingham Bridge tide gauge; Manly Hydraulics Laboratory), providing suitable conditions conducive for wetland inundation and immature mosquito populations during each sampling period.

## Results

A total of 16,467 mosquitoes was collected at all sites across the two sampling periods representing seven genera and 22 species (Table 1). The most commonly collected mosquitoes were *Ve. funerea* (41.92% of total mosquitoes collected), *Ae. vigilax* (30.23% of total mosquitoes collected), *Cx. sitiens* (12.97% of total mosquitoes collected), *Cx. annulirostris* (5.73% of total mosquitoes collected), and *Ae. notoscriptus* (2.96% of total mosquitoes collected). The relative abundance and diversity of mosquito collections at each site across the two sampling periods remained relatively comparable. During January 2021, a total of 9,181 mosquitoes was collected while during February 2021, a total of 7,286 mosquitoes was collected. The mosquito fauna collected at the Ballina Heights reference site was reflective of its location beyond the flight range of mosquitoes associated with estuarine wetlands and the concomitant mosquito species recorded were those typically associated with freshwater habitats with a relatively greater abundance of species that included *Aedes multiplex* (Diptera: Culicidae) and *Ae. procax*, and *Cq. linealis* (Webb et al. 2016).

**Table 1.** Mean trap night abundance of mosquito species collected as adults at five sampling locations, on two occasions, January 2021 and February 2021. Ballina, NSW, Australia.

| Species/Trap Site | River Oaks Estate<br>Conservation Area<br>(trap nights=8) |  | North Creek<br>(trap nights =4) |  | Little Fishery Creek<br>(trap nights =4) |  | Fishery Creek<br>(trap nights =4) |  | Ballina Heights<br>(trap nights =2) |  |
| --- | --- | --- | --- | --- | --- | --- | --- | --- | --- | --- |
|  | January<br>2021 | February<br>2021 | January<br>2021 | February<br>2021 | January<br>2021 | February<br>2021 | January<br>2021 | February<br>2021 | January<br>2021 | February<br>2021 |
| *Aedes alternans* | 1.3 | 1.2 | 12.5 | 9.0 | 2.3 | 3.0 | 3.0 | 3.5 | 1.0 | 1.0 |
| *Aedes flavifrons* | 0.0 | 0.0 | 1.0 | 0.0 | 0.0 | 0.0 | 0.0 | 0.0 | 0.0 | 0.0 |
| *Aedes kochi* | 2.0 | 0.0 | 1.0 | 2.0 | 0.0 | 0.0 | 0.0 | 0.0 | 0.0 | 2.5 |
| *Aedes multiplex* | 3.6 | 0.0 | 9.5 | 9.0 | 8.3 | 1.5 | 1.8 | 0.0 | 67.5 | 32.5 |
| *Aedes notoscriptus* | 15.5 | 8.1 | 13.8 | 8.8 | 13.3 | 4.0 | 7.0 | 4.5 | 29.5 | 24.5 |
| *Aedes procax* | 2.5 | 1.0 | 4.3 | 0.0 | 2.0 | 0.0 | 0.0 | 0.0 | 10.0 | 23.0 |
| *Aedes vigilax* | 31.4 | 28.5 | 634.3 | 310.5 | 85.5 | 37.5 | 15.8 | 33.5 | 6.0 | 9.5 |
| *Aedes vittiger* | 1.0 | 0.0 | 0.0 | 0.0 | 0.0 | 0.0 | 1.0 | 0.0 | 0.0 | 0.0 |
| *Anopheles annulipes* | 1.0 | 0.0 | 2.0 | 2.3 | 1.0 | 0.0 | 0.0 | 0.0 | 1.0 | 0.0 |
| *Coquillettidia linealis* | 0.0 | 8.0 | 5.5 | 31.5 | 0.0 | 2.0 | 0.0 | 2.0 | 14.0 | 51.5 |
| *Coquillettidia variagata* | 1.0 | 0.0 | 0.0 | 2.0 | 0.0 | 0.0 | 0.0 | 0.0 | 0.0 | 3.0 |
| *Coquillettidia xanthogaster* | 1.3 | 5.0 | 8.8 | 32.8 | 1.5 | 8.0 | 3.8 | 2.3 | 17.0 | 3.5 |
| *Culex annulirostris* | 13.0 | 14.3 | 25.8 | 85.5 | 5.3 | 27.5 | 2.5 | 21.7 | 36.5 | 29.0 |
| *Culex bitaeniorhynchus* | 1.0 | 0.0 | 0.0 | 0.0 | 2.0 | 2.0 | 1.0 | 2.0 | 3.0 | 1.0 |
| *Culex pipiens. molestus* | 0.0 | 0.0 | 0.0 | 0.0 | 1.0 | 0.0 | 0.0 | 0.0 | 0.0 | 0.0 |
| *Culex orbostiensis* | 5.5 | 0.0 | 6.8 | 6.3 | 2.5 | 0.0 | 0.0 | 1.0 | 49.0 | 6.0 |
| *Culex quiquefasciatus* | 13.5 | 2.3 | 1.0 | 2.0 | 16.5 | 21.7 | 25.0 | 4.5 | 0.0 | 2.0 |
| *Culex sitiens* | 39.1 | 20.8 | 81.8 | 120.0 | 81.7 | 74.0 | 41.5 | 28.0 | 7.5 | 8.0 |
| *Mansonia uniformis* | 0.0 | 1.2 | 1.0 | 12.0 | 0.0 | 2.0 | 0.0 | 0.0 | 0.0 | 14.5 |
| *Uranotaenia lateralis* | 0.0 | 0.0 | 0.0 | 0.0 | 0.0 | 0.0 | 0.0 | 0.0 | 1.0 | 1.0 |
| *Verrallina funerea* | 25.9 | 23.6 | 714.8 | 720.5 | 29.5 | 43.8 | 21.8 | 47.0 | 104.0 | 8.0 |
| *Verrallina* Marks 52 | 0.0 | 0.0 | 0.0 | 0.0 | 0.0 | 0.0 | 0.0 | 0.0 | 10.5 | 1.0 |
| **TOTAL** | **131.6** | **102.0** | **1512.3** | **1539.0** | **222.3** | **213.8** | **119.5** | **140.3** | **356.0** | **218.5** |

## Discussion

The key mosquitoes of pest and public health concern identified in this study were *Ae. vigilax* and *Ve. funerea*, known to be nuisance biting pests and population abundance has identified as a contributing factor to outbreaks of mosquito-borne disease (Ritchie et al. 1997; Ryan et al. 1999; Jeffery et al. 2006; Jacups et al. 2008). The relative abundance of both species was comparable between the estuarine wetland sites of River Oaks Estate Conservation Area, Fishery Creek, and Little Fishery Creek but substantially lower compared North Creek where the highest abundance of both species was recorded. The results of this investigation demonstrate that while suitable habitats these mosquitoes are present throughout the wider estuary, their productivity may not be directly comparable. This phenomenon has been observed in other estuarine wetlands where the abundance of *Ae. vigilax* exhibits a high degree of spatial variability despite extensive estuarine wetlands that represent suitable habitats (Claflin and Webb 2016) or where the temporal differences in brackish water habitat productivity of *Ve. funerea* varies depending on either rainfall or tidal inundation of habitats (Jeffery et al. 2005).

When assessing the local threats posed by these mosquitoes, it is important to consider both the productivity of local habitats as well as the dispersal of these mosquitoes. It is noteworthy that there are distinct differences in the documented dispersal ranges of these two mosquitoes with *Ae. vigilax* dispersing many kilometres from larval habitats (Webb and Russell 2019; Johnson et al. 2020) while *Ve. funearea* is thought to travel only short distances away from their coastal swamp forests and she-oak woodland habitats (Ryan et al. 1999; Webb et al. 2016). This difference in dispersal has implications for mosquito control or other integrated mosquito management programs. While there are ecologically sustainable control measures available for both species (Russell and Kay 2008; Ryan and Kay 2000; Jeffery et al. 2014), prioritising areas for control may require an assessment of residential and/or recreational areas at greatest risk and, similarly, consideration of current and future mosquito impacts may need to be considered when assessing various land uses within areas prone to potentially substantial mosquito impacts.

Additional consideration should be given to those mosquitoes found in urban areas adjacent to natural environments. In residential and industrial areas, mosquitoes associated with water-holding containers, such as *Ae. notoscriptus*, may also be a concern given it is one of the most widespread nuisance-biting mosquitoes, given its close association with water-holding containers in urban settings, and potential vector of arboviruses in Australia (Watson and Kay 1998; Kay et al. 2008). While this mosquito may not disperse more than 200m from larval habitats, where urban areas are adjacent to wetlands or other natural areas, there may be some confusion related to the source of mosquito species driving nuisance impacts. There are other mosquitoes found in urban habitats that may cause nuisance problems. *Culex quinquefasciatus* (Diptera: Culicidae) and *Culex pipiens molestus* (Diptera: Culicidae) were both recorded in this investigation and, although the numbers were collected were very low, they may be a source of pest problems under some circumstances.

Mosquitoes are a natural part of coastal wetlands in the Northern River region of NSW but the results of this investigation highlight the spatial and, potentially, temporal differences, in abundance and diversity. When authorities consider the most appropriate strategies to manage the pest and public health risks associated with mosquitoes, consideration needs to be given to site-specific mosquito risk assessment as opposed to any generalised risk assessment based on vegetation communities alone. Informing such management decisions with mosquito population data will assist the effectiveness of management programs.

## Acknowledgements

The authors with that thank Kristy Bell and Ian Gaskell from Ballina Shire Council for assistance in providing background information on local wetlands and mosquito populations that assisted the design of methodology and interpretation of results presented here. We acknowledge the Bundjalung people, the traditional owners of the land on which this research was conducted.

